# Cyto-nuclear linkage disequilibrium resulting from admixture

**DOI:** 10.1101/2020.11.24.396903

**Authors:** Peter D. Fields, David E. McCauley, Douglas R. Taylor

**Affiliations:** Department of Biology, University of Virginia, PO Box 400328, Charlottesville, VA 22904-4328; University of Basel, Zoological Institute, Vesalgasse 1, Basel, CH-4051, Switzerland; Department of Biological Sciences, Vanderbilt University, Nashville, TN 37235

## Abstract

Previous studies of North American populations of the invasive plant *Silene latifolia* showed significant cyto-nuclear linkage disequilibrium (CNLD) between SNP variants of a mitochondrial gene (*atp*1) and the most common allele at nuclear microsatellite loci. Fields *et al*. (2014) hypothesized that this CNLD arose partially as a consequence of admixture that occurred during the colonization of North American (NA) populations of *S latifolia* via seed dispersal from genetically differentiated European populations that represent a portion of the native range of this species. In order to evaluate the plausibility of the admixture hypothesis, as opposed to metapopulation processes alone, we estimated CNLD for these same loci using data collected from eastern (EEU) and western (WEU) European populations of *S. latifolia* known to be genetically differentiated and likely sources of the spread of the study species to North America. We show that the CNLD found previously in NA populations of *S. latifolia* can be attributed to admixture of the previously isolated European demes coupled with decay since that time. Our applied framework allows the separation of the forces generating and dissolving statistical associations between alleles in cytoplasmic organelles and the nuclear genome and may thus be of utility in the study of plant or animal microbiomes.

## INTRODUCTION

The magnitude and distribution of genetic variation across the recently established range of an invasive species is a reflection of the demographic history of demes from the native range, colonization events, and other evolutionary processes such as natural selection and genetic drift within the introduced range (Taylor & Keller, 2007, Dlugosch & Parker, 2008, Hufbauer, 2008, Calsbeek et al., 2011, Colautti & Barrett, 2011, Keller et al., 2014). For our purposes genetic variation can be defined as the degree of polymorphism at some number of loci as measured in aggregate across the new range, or as measured both within local populations as well as among those populations (i.e., considering the geographic population genetic structure). Measurements of variation could be extended beyond the standard locus-by-locus variety to include the distribution within and among populations of various multi-locus genotypes. These could include evaluation of non-random associations of alleles across loci (linkage disequilibrium or LD), again considering the genetic composition of all populations in aggregate or on a population-by-population basis.

In order to understand the determinants of genetic variation described above, it would be useful to consider mechanisms of short and long-range gene dispersal, along with the influence of the mating system, on multi-locus genotypic diversity. Note that in many species of seed plants most genes have the potential to disperse in both seeds and pollen. However, because the chloroplast and mitochondrial genomes tend to be extensively (or entirely) maternally inherited (McCauley, 2013) the movement of pollen would appear to be a more important mechanism for long distance dispersal of nuclear genes as compared to that of organellar genes. This distinction would appear to be especially relevant to the emigration of genetic diversity during the early stages of an invasion since a new population cannot be founded initially by pollen alone and thus requires seed dispersal (Baker & Stebbins, 1965). On the other hand, long distance seed dispersal can transport cyto-nuclear genotype combinations that exist in the source populations (Hu, 2008). That is, when cyto-nuclear linkage disequilibrium (CNLD) is found in an invasive plant species one must consider whether that CNLD is a reflection of the genetic composition of the populations from which colonists were drawn. The possibilities include that the CNLD occurred in some populations in the native range, or that it was created as colonists from genetically differentiated populations in the native range underwent admixture at or after the time of the invasion. However, it is known from theory that CNLD found in neutral variation should decline by 50% per generation in panmictic populations so CNLD persistence is problematic whatever the origin (Fields et al., 2014, Hu, 2008). Note also, though, that numerous species of plants are predominant selfers, and that recurrent self-fertilization limits the opportunity for the mixing of alleles between genomes. In addition to helping to reveal the history of populations regarding patterns of gene flow and mating system, study of CNLD can also be useful when considering the role of cyto-nuclear co-evolution, including selective phenomenon leading to coadaptation or conflict between the genomes (Fields et al., 2014, Sloan, 2015, Bock et al., 2014) and the transfer of genetic information between genomes (Sloan, 2015).

In a recent study of the plant *Silene latifolia*, Fields et al. (2014) identified high levels of cyto-nuclear linkage disequilibrium (CNLD) between variants of a SNP that occurs within the mitochondrial gene *atp1* and alleles at a number of nuclear microsatellite loci. Samples were taken from populations located in southwestern Virginia, USA, within what is assumed to be the eastern portion of the invasive range of *S. latifolia* in North America (Keller et al., 2012). A comparison to genotypes found in the same *S. latifolia* populations sampled 20 years (∼ 7 generations) prior to the more recent samples revealed only modest change in mean CNLD through time. Fields et al. (2014) speculated that the CNLD that was observed could be traced to1) metapopulation dynamics, in particular founder effects associated with a propagule pool (Slatkin, 1977, Wade & McCauley, 1988, Whitlock & McCauley, 1990) colonization dynamic, and 2) the colonization process by which the European native *S. latifolia* became established in this region of the eastern U.S. via seed dispersal. The relative temporal stability of the observed CNLD could be due to the fact that *S. latifolia* displays a high degree of local population structure in this area (McCauley, 1994) and that this would limit opportunities for the outbreeding needed for the erosion of any CNLD that arose when those populations were established. A link between CNLD and the establishment of populations would be especially likely if the various European populations contributing colonists (either directly or indirectly via dispersal from other recently established North American populations) displayed some degree of cyto-nuclear differentiation that could contribute to CNLD, especially with admixture.

Extensive recent sampling across much of the putative native range of *S. latifolia* in Europe has revealed geographic genetic structuring that is especially noticeable when populations from eastern and western Europe are compared (Keller et al., 2012, Keller et al., 2014, Taylor & Keller, 2007). Here we investigate the plausibility that the CNLD in North American populations found by Fields et al. (2014) could be a consequence of the recent establishment of *S. latifolia* in North America and admixture between divergent European populations during hypothetical invasion events. We find that different invasion scenarios, combined with different forms of cyto-nuclear population differentiation in the native range, could have enhanced or diminished the CNLD observed in the invasive range. Application of cyto-nuclear genotyping, in addition to nuclear genotyping alone, provides additional insights for the origin of statistical associations of genetic variation among genomic regions, and the relative roles of historical demography and ongoing metapopulation processes.

### Theoretical expectations for shifts in CNLD

Consider a population of individuals that are characterized at two loci, one cytoplasmic (*C*) and one nuclear (*N*) with two alleles at each locus (*C*_1_ & *C*_2_; *N*_1_ & *N*_2_). This population could be described according to the frequency (*p*) of one of the alleles at each locus (i.e. *p*_*C,1*_ and *p*_*N,1*_) and by the frequency of the cytonuclear genotypes (i.e. *p*_*C,N,1 etc*_). The disequilibrium between *C*_1_ & *N*_1_, *D*_*C-N*_, could be defined in this population as *D*_*C-N*_ = [*p*_*C,N,1*_ – ((*p*_*C,1*_)(*p*_*N,1*_))]. Thus, *D*_*C-N*_ determines whether two alleles are associated within individuals, more (positive) or less (negative) often than random and is mathematically equivalent to the covariance between alleles. Note that the equation for LD is closely related to a covariance (symbolized as *D*) with maximum values constrained by the gene frequencies in the sample. Another standardized measure of CNLD (*D’*) is also commonly reported (e.g. in Fields et al. (2014)). This standardization constrains values of *D’* to fall between −1 and 1 (negative and perfect association of alleles, respectively).

Now consider *p*_*C,1*_, *p*_*N,1*_ and *p*_*C,N,1*_ for each of *k* = 2 or more populations individually. From this information, the overall CNLD value could be calculated in two ways. First, *D*_*C-N*_ can be calculated for each of the *k* populations, and assuming the populations are identical in size, these values could be averaged to obtain 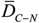. Alternatively, individual genotypes could be pooled across the *k* populations and used to calculate values of *p*_*C,1*_, *p*_*N,1*,_, and *p*_*C,N,1*_ *p*_*C,1*_*p*_*N,1*_ and 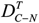 based on the pooled total set of individual genotypes.

Cytonuclear disequilibrium, and LD in general, may arise from several sources or by processes operating at different spatial scales. First, at a larger spatial scale, CNLD will arise naturally as a result of population genetic divergence at both nuclear and cytoplasmic loci. Individual populations may also display variation in the population-specific values of *D*_C-N_ according to how the alleles at the nuclear and cytoplasmic loci are organized into two-locus genotypes in each population, which could be a function of a variety of forces operating within populations (e.g. mating structure, demographic history, selection, etc.).

One might predict how the cyto-nuclear genetic structure of a set of *k* populations determines **D**^**T**^_**C-N**_ in a population originating from mixing individuals from those populations. It can be shown that

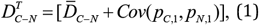

where *Cov*(*p*_*C*,1_, *p*_*N*, 1_) is the covariance of the *k* population-specific cytoplasmic and nuclear allele frequencies across populations contributing equally to the mixture (p. 550, Hedrick, 2011). Note, this equation does not apply directly to *D’*.

Figure 2 illustrates the application of this equation to several hypothetical sets of *k* = 2 source populations. From this equation it can be seen that mixing individuals from more than one source can result in a *D*_*T*C-N_ in the new population higher than the average *D*^*T*^_C-N_ in the source populations if *Cov*(*p*_*C*,1_, *p*_*N*, 1_) > 0 is (Figure 2, scenario 1). *Cov*(*p*_*C*,1_, *p*_*N*, 1_) would be positive if certain combinations of cytoplasmic and nuclear alleles coexist in the same populations more often than a random association across populations. Mixing can diminish *D*_C-N_ if *Cov*(*p*_*C*,1_, *p*_*N*, 1_) is negative (Figure 2, scenario 2; scenario 3), as would be the case if certain cyto-nuclear alleles co-occur less often than random in the source populations. It can have no effect on *D*^*T*^_C-N_ as compared to average population-specific CNLD (ignoring stochastic sampling) if *Cov*(*p*_*C*,1_, *p*_*N*, 1_) = 0, as (Figure 2 scenario 4; scenario 5). *Cov*(*p*_*C*,1_, *p*_*N*, 1_) = 0 would be the case if the nature of cyto-nuclear associations is consistent across populations.

### CNLD in the European range of S. latifolia

Samples of *S. latifolia* were collected from 85 individuals distributed across 12 populations in the species’ native European range (Figure 1), a subset of the total sample used by Keller et al. (2012). Genomic DNA was extracted as described in Keller et al. (2012) using methods adapted from Edwards et al. (1991) and Slotta et al. (2008). Individuals were genotyped at 14 microsatellite markers derived from *S. latifolia* (slat18, slat32, slat33, slat48, slat72, slat85) described by Molecular Ecology Resources Primer Development et al. (2010), and (SL_eSSR04, SL_eSSR06, SL_eSSR09, SL_eSSR12, SL_eSS16, SL_eSSR20, SL_eSSR29, SL_eSSR30) described by Moccia et al. (2009). Fragment analysis was conducted using an ABI 313xl automated sequencer, with genotypes determined using GENEMAPPER v.3.0 software (Applied Biosystems) and binned using the program TANDEM (Matschiner & Salzburger, 2009).

**Figure 1.**
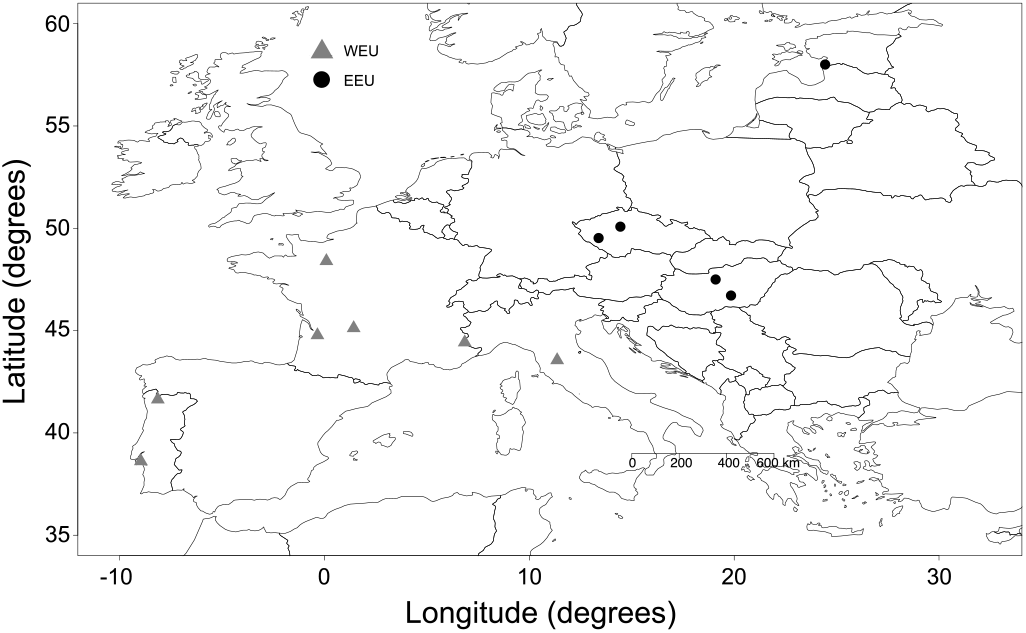
European sampling localities. Grey dots indicate samples from the Western European (WEU) deme and black dots indicate samples from Eastern European (EEU) deme.

**Figure 2.**
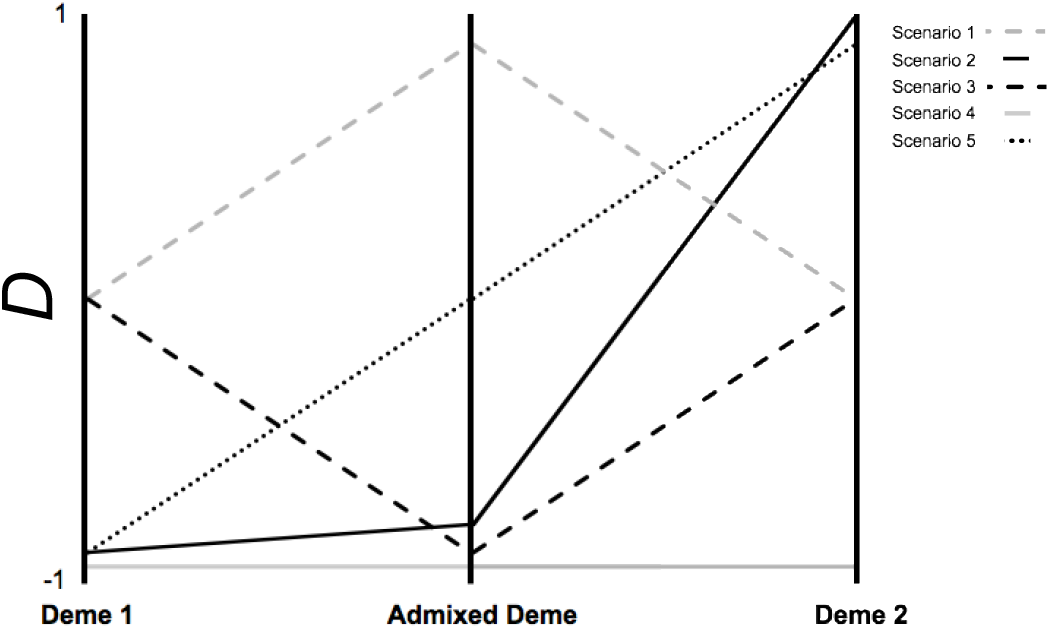
Hypothetical consequences for CNLD *D* values when populations that differ in their respective values of *D*, due to variable genetic compositions, contribute individuals to form an admixed deme. Five types of mixtures/scenarios are described in the text.

We followed the methods described by Keller et al. (2012) in order to distinguish the two distinct, post-glacial demes in *S. latifolia*’s native range. Briefly, individuals are assigned to Eastern and Western Europe (EEU and WEU, respectively) based upon a suture zone running north from the Alps to the North Sea, and has been described previously as different routes of expansion for *S. latifolia* out of distinct Mediterranean refugia following Holocene warming (Hewitt, 1996, Petit et al., 2003, Taylor & Keller, 2007, Keller et al., 2009, Keller et al., 2012). Using these *a priori* expectations, Keller et al. (2012) applied Bayesian clustering using the program STRUCTURE v.2.2 (Pritchard et al., 2000, Falush et al., 2003), assuming an admixture model with allele frequencies correlated and remaining parameters set as a default over 10 independent runs with 200,000 MCMC iterations and a burn-in period of 50,000 iterations. Here we utilize cluster assignments assuming K=2 representing EEU and WEU, respectively (Keller et al., 2012, Taylor & Keller, 2007).

In order to estimate CNLD, we followed the approach of Fields et al. (2014). The same genomic samples used for the microsatellite analysis were assayed for a SNP in the mitochondrial gene *atp1* using a PCR/RFLP method (McCauley & Ellis, 2008, Fields et al., 2014). PCR products were digested with *AluI* (New England Biolabs), electrophoresed on 4% Metaphor agarose gel (Lonza Inc, Rockland, ME, USA), and stained with ethidium bromide in order to resolve the two possible alleles.

We calculated the observed (*H*_O_) and expected (*H*_E_) heterozygosity of our nuclear genetic markers using the software GenoDive version 2.0b21 (Meirmans & Van Tienderen, 2004). Estimates of genetic substructure using hierarchical F-statistics were calculated using the software FSTAT 2.9.3.2 (Goudet, 2002), with significant deviations from panmixia assessed by testing for Hardy–Weinberg Equilibrium with 10,000 permutations and *α* =0.05.

We estimated nuclear-nuclear LD (hereafter, nuclear LD) following the methods described in Fields et al. (2014). Statistical significance of nuclear LD between pairs of loci for post-glacial demes, as well as within the full European sample, was determined using a Monte-Carlo approximation of Fisher’s exact test implemented in the software Arlequin (Excoffier & Lischer, 2010), which assumes a null hypothesis of random allelic assortment. Arlequin uses a Markov chain extension of Fisher’s exact test for *RxC* contingency tables (Slatkin, 1994, Li & Merila, 2010). 100,000 alternative tables were explored by the Markov chain (Slate & Pemberton, 2007, Li & Merila, 2010).

We follow the methods described in Fields et al. (2014) to estimate cyto-nuclear *D* and *D’*. Briefly, we followed the approach of Basten and Asmussen (1997), using the program CNDm to estimate a standardized estimate an allelic *D* and *D’* between each nuclear locus and atp1 mtDNA locus. CNDm uses a Monte-Carlo approach to approximate Fisher’s exact test for RxC contingency tables and tests for significant deviations from the null hypothesis of no allelic association (Basten & Asmussen, 1997). For this analysis, all nuclear loci were treated as bi-allelic by pooling all alleles other than the most common allele in the full European sample into a single composite allele (as in Latta et al., 2001).

## RESULTS AND DISCUSSION

Cyto-nuclear genotypes were obtained for a total of 85 European (EU) individuals (52 individuals from the Western European Deme (WEU) and 33 individuals from the Eastern European Deme (EEU)). For the *atp1* SNP the most common variant over the full European range occurred at a frequency of 0.535 (0.929 in EEU, and 0.254 in WEU). Table 1 presents a summary of global estimates of nuclear genetic diversity, population substructure (*F*_IT_) and a summary of nuclear LD for the EEU, WEU, and full EU sample. All microsatellite loci were polymorphic (> 1 allele) in both regions, and ranged between 2 and 21 alleles. As in Fields et al. (2014), individual loci showed a large range of *H*_O_ (0.061-0.761, mean = 0.38) and *H*_E_ (0.134-0.902, mean = 0.63). Average *F*_IT_ for the whole EU sample was 0.498 (0.17-0.94), with all loci exhibiting significant (*P* < 0.05) deviations from panmixia, corroborating deviations between *H*_O_ and *H*_E_. The observed *F*_IT_ was larger than that reported in Fields et al. (2014), which is to be expected given differences in scale and the opportunity for admixture that took place during the invasion of the North American continent (Keller et al., 2012). The average number of loci pairs showing significant nuclear-nuclear LD was approximately equal in WEU and EEU (3.86 and 5.71, respectively), though the full sample showed approximately twice as much (9.86). Table 2 presents the observed distribution of *D* and *D’* values for EEU, WEU, and the full EU sample, respectively, for each of 14 nuclear microsatellites and the mitochondrial *atp1* SNP. Similarly to Fields et al. (2014), overall mean levels of LD were consistent, while specific locus values varied widely.

**Table 1.**
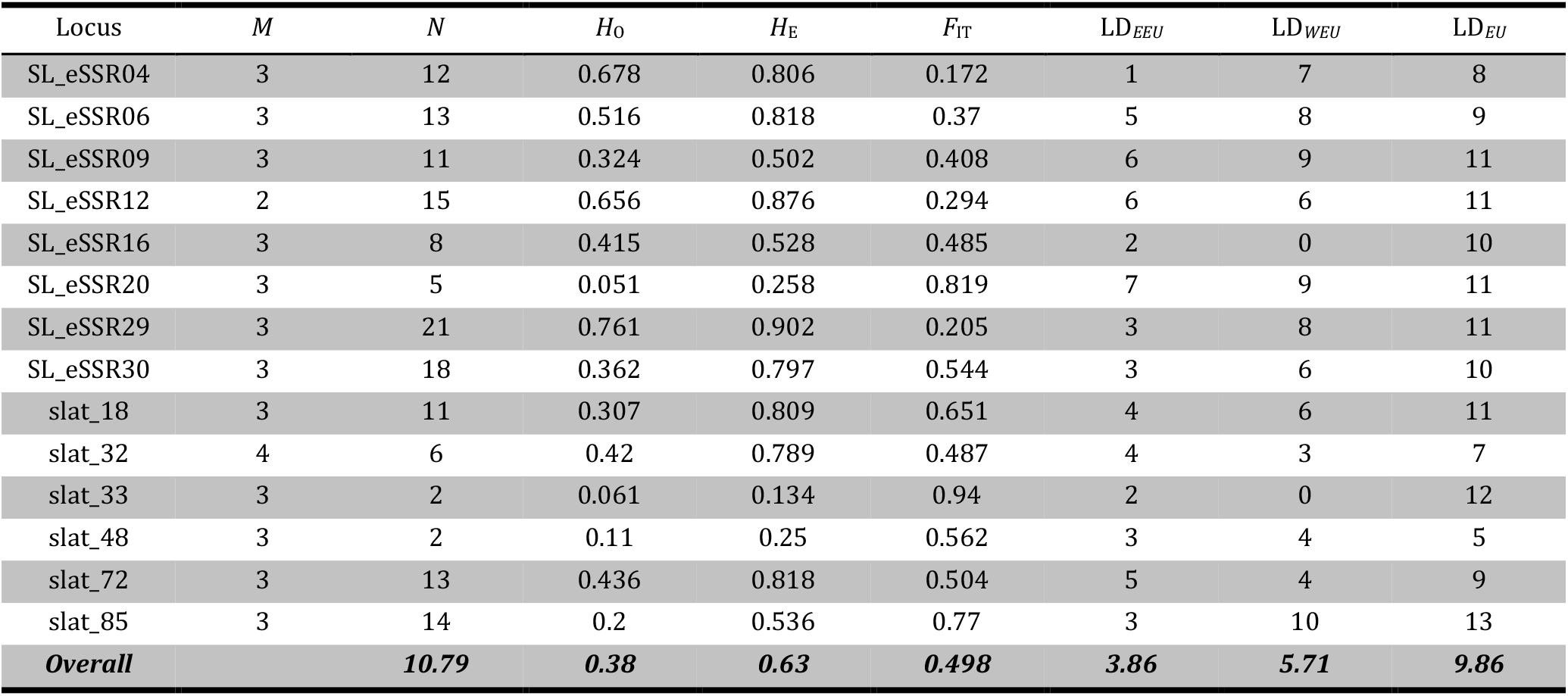
Global estimates of genetic diversity, population substructure, and linkage disequilibrium across Europe. *M*=repeat size of the marker; *N*=number of observed alleles; *P*=allele frequency of the most frequent allele. *H*_O_ and *H*_E_ are observed and expected heterozygosity, respectively, of each sampled microsatellite locus; *F*_IT_ is the correlation of alleles within individuals relative to the continental sample as a whole, and can be interpreted as an approximate measure of deviation from panmixia. Statistical deviations from panmixia are indicated by the symbols *P<0.05, **P<0.01 and ***P<0.001) and LD_*EEU*_ is an indication of the number of other nuclear loci for which each focal nuclear locus exhibits a non-random association in the EEU deme; LD_*WEU*_ is an indication of the number of other nuclear loci for which each focal nuclear locus exhibits a non-random association in the WEU deme; and LD_*EU*_ is an indication of the number of other nuclear loci for which each focal nuclear locus exhibits a non-random association across the whole European sample, based upon an approximation of Fisher’s exact test of random association and a *P*-value ≤ 0.05.

| Locus | $M$ | $N$ | $H_0$ | $H_E$ | $F_{IT}$ | $LD_{EEU}$ | $LD_{WEU}$ | $LD_{EU}$ |
| --- | --- | --- | --- | --- | --- | --- | --- | --- |
| SL_eSSR04 | 3 | 12 | 0.678 | 0.806 | 0.172 | 1 | 7 | 8 |
| SL_eSSR06 | 3 | 13 | 0.516 | 0.818 | 0.37 | 5 | 8 | 9 |
| SL_eSSR09 | 3 | 11 | 0.324 | 0.502 | 0.408 | 6 | 9 | 11 |
| SL_eSSR12 | 2 | 15 | 0.656 | 0.876 | 0.294 | 6 | 6 | 11 |
| SL_eSSR16 | 3 | 8 | 0.415 | 0.528 | 0.485 | 2 | 0 | 10 |
| SL_eSSR20 | 3 | 5 | 0.051 | 0.258 | 0.819 | 7 | 9 | 11 |
| SL_eSSR29 | 3 | 21 | 0.761 | 0.902 | 0.205 | 3 | 8 | 11 |
| SL_eSSR30 | 3 | 18 | 0.362 | 0.797 | 0.544 | 3 | 6 | 10 |
| slat_18 | 3 | 11 | 0.307 | 0.809 | 0.651 | 4 | 6 | 11 |
| slat_32 | 4 | 6 | 0.42 | 0.789 | 0.487 | 4 | 3 | 7 |
| slat_33 | 3 | 2 | 0.061 | 0.134 | 0.94 | 2 | 0 | 12 |
| slat_48 | 3 | 2 | 0.11 | 0.25 | 0.562 | 3 | 4 | 5 |
| slat_72 | 3 | 13 | 0.436 | 0.818 | 0.504 | 5 | 4 | 9 |
| slat_85 | 3 | 14 | 0.2 | 0.536 | 0.77 | 3 | 10 | 13 |
| **Overall** |  | **10.79** | **0.38** | **0.63** | **0.498** | **3.86** | **5.71** | **9.86** |

**Table 2.** Values of Cyto-nuclear linkage disequilibrium (D or D’) between the mitochondrial ATP1 SNP and the most common allele (at frequency p) of each of 14 nuclear microsatellite markers calculated from sample taken at two localities in Europe (EEU & WEU) and a mixture of those European samples (EU). Also presented are D and D” calculated from North American (NA) samples reported by Fields *et al*. (2014).

| Locus | EEU |  |  | WEU |  |  | EU |  |  | NA |  |  |
| --- | --- | --- | --- | --- | --- | --- | --- | --- | --- | --- | --- | --- |
|  | P | D | D' | P | D | D' | P | D | D' | P | D | D' |
| SL_eSSR04 | 0.329 | 0 | 0 | 0.237 | -0.0288 | -0.25 | 0.276 | -0.0099 | -0.0682 | 0.498 | 0.0255 | 0.1737* |
| SL_eSSR06 | 0.268 | -0.0069 | -0.061 | 0.395 | 0.0481 | 0.25 | 0.342 | 0.0137 | 0.1146*** | 0.262 | -0.0257 | -0.1751** |
| SL_eSSR09 | 0.744 | 0.0234 | 0.2361 | 0.623 | -0.1202 | -0.641 | 0.672 | -0.0579 | -0.4504** | 0.743 | 0.0312 | 0.2161** |
| SL_eSSR12 | 0.232 | -0.0386 | -0.3889 | 0.061 | 0.0337 | 1 | 0.133 | 0.0223 | 0.387*** | 0.315 | 0.0448 | 0.3243*** |
| SL_eSSR16 | 0.207 | -0.022 | -0.2254 | 0.816 | 0.0048 | 0.0526 | 0.561 | -0.0513 | -0.3266*** | 0.544 | 0.0539 | 0.3258*** |
| SL_eSSR20 | 0.902 | -0.011 | -1 | 0.798 | -0.0481 | -0.5 | 0.842 | -0.0237 | -0.4466*** | 0.953 | -0.0078 | -0.3721 |
| SL_eSSR29 | 0.25 | -0.0413 | -0.4762 | 0.026 | -0.0144 | -1 | 0.119 | -0.0064 | -0.0836*** | 0.203 | 0.0254 | 0.2841** |
| SL_eSSR30 | 0.427 | 0.0317 | 0.4107 | 0.298 | -0.024 | -0.1515 | 0.352 | 0.0057 | 0.042 | 0.389 | 0.0117 | 0.0691 |
| slat_18 | 0.038 | -0.022 | -0.5926 | 0.429 | -0.0481 | -0.25 | 0.268 | -0.0636 | -0.4032*** | 0.402 | 0.0125 | 0.071 |
| slat_32 | 0.351 | 0.0069 | 0.1316 | 0.278 | 0.0288 | 0.2143 | 0.308 | 0.0217 | 0.2088 | 0.316 | -0.0561 | -0.3161*** |
| slat_33 | 0.11 | -0.0262 | -0.3016 | 0.965 | 0 | 0 | 0.607 | -0.0748 | -0.5362*** | 0.607 | 0.1042 | 0.4868*** |
| slat_48 | 0.808 | 0.0096 | 0.0833 | 0.9 | 0.0337 | 0.6364 | 0.862 | 0.0174 | 0.1981 | 0.582 | 0.0475 | 0.2415*** |
| slat_72 | 0.325 | 0.0193 | 0.3889 | 0.219 | 0.0673 | 0.5833 | 0.263 | 0.0518 | 0.5573* | 0.308 | 0.0026 | 0.0193 |
| slat_85 | 0.878 | 0.0358 | 0.3714 | 0.145 | -0.0577 | -0.75 | 0.458 | 0.0322 | 0.1995*** | 0.516 | -0.0256 | -0.1492* |

The main goal of this study was to ask whether the levels of CNLD seen by Fields et al (2014) in a North American (NA) sample of *S. latifolia* could plausibly have resulted from admixture of colonists from the European range of this species, Given this, it would seem useful to compare values of D and/or D’ obtained from North America (NA) to those from Europe. To that end we have included the pooled values of *D’* from Table 2 of Fields *et al*. (2014) in their Table 2, along with associated values of *D* calculated from the same data but not presented in Fields et al. (2014).

From our Table 2 it can be seen that the range of *D’* for NA was -.372 to.487 (mean = .086). For EU the range was -.536 .to .557 (mean = .043). In a similar comparison of *D* values the range for NA was -.078 to .1042 (mean = .016). For EU the range was -.075 to 0.52 (mean = -.009)

Given a fair degree of locus-to-locus variation in the nuclear locus-specific values of *D*’ and *D* in EU and NA, an additional method a comparing the similarity of CNLD in EU and NA would be to examine the consistency of these locus-specific values by calculating their product-moment correlation (*r*). That correlation is *r* = -.208 for *D*’ and *r* = −0.397 for *D*. Neither *r* value is significantly different from zero under a null hypothesis of *r* = 0 (p > 0.05, df = 12 in each case), suggesting drift-like dynamics are responsible for shifting CNLD rather than selective agents.

What is the impact of the covariance of cyto-nuclear allele frequencies across EEU and WEU on EU *D* values? Recall from eq. 1 and Fig 1 that represent various “scenarios” that such covariance can either increase or decrease the magnitude of EU D relative to the simple mean of site-specific D values. Inspection of Fig. 3A suggests that all of the scenarios are approximated, depending on the nuclear locus in question. Locus-specific Shifts in CNLD D values range from a shift of .058 (locus SLeSSR12) to a decrease of -.1062 (locus Slat 33) (mean shift in D = -.002).

**Figure 3.**
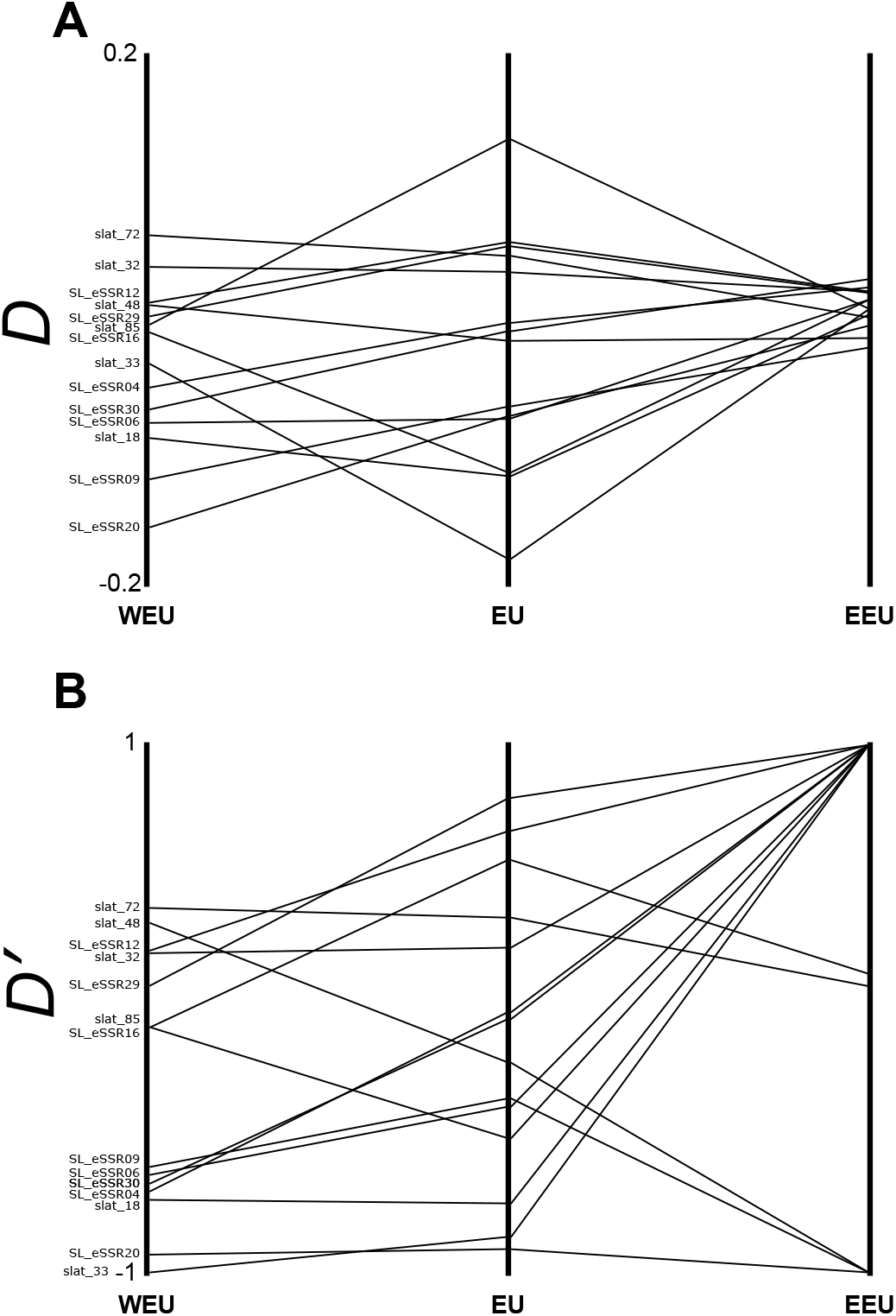
Locus by locus CNLD Values of *D* (3A) or *D*’ (3B) obtained from single European sources (EEU and WEU) and from pooling samples from those sources (EU). The shift predicted from eq. 1 can be visualized by noting that the D value seen in EU are not always the average of the locus-specific D found in EEU and WEU.

Examination of Table 2 and Figure 3A suggests a large role of differing frequency of the most common cytoplasmic allele, wherein the greater frequency of the allele in EEU leads to low values of *D* for most loci, and a larger range of *D* in WEU. Figure 3B describes a similar pattern for *D*’, wherein the near fixation of one cytoplasmic allele in the EEU deme generates a very strong signal of CNLD across 12 of 14 nuclear loci. However, both Figure 3A and 3B imply just how drastically admixture of two highly divergent demes can shift the overall mean and variance of CNLD. Thus, our results are in least partial support of our previous hypothesis that the CNLD found in our North American *S. latifolia* samples could result, in part, from admixture of colonists from two or more sources in the native range of *S. latifolia* that contributed to the expansion of the range of this species into eastern North America.

## CONCLUSION

Linkage disequilibrium, either nuclear-nuclear or cyto-nuclear, provides a useful summary of the demographic processes taking place within and among species. While estimation of nuclear-nuclear LD is common, estimation of CNLD is much less so. This is at least in part due to a much less extensive exploration, both theoretical and empirically, of CNLD. This state seems particularly problematic given the general realization of the role of cyto-nuclear interactions in important evolutionary processes, including local adaptation (e.g. Bock et al., 2014) and speciation (e.g. Trier et al., 2014), as well as co-evolutionary dynamics more generally (Sloan, 2015). Fields et al. (2014) provided a temporal study of the how CNLD will shift over time as a result of metapopulation processes. The present study shows that the incomplete admixture of *S. latifolia* in North America also contributed to CNLD, as well as outlining how allele frequencies in the species native range resolved into present patterns of CNLD in the invaded range. Application of the presented framework, and modifications therein, should allow for a fuller understanding the role of selective and non-selective forces in generating patterns of LD both within and among genomes.

